# Measuring aptamer folding energy using a molecular clamp

**DOI:** 10.1101/2020.02.10.941203

**Authors:** Hao Qu, Qihui Ma, Lu Wang, Yu Mao, Michael Eisenstein, Hyongsok Tom Soh, Lei Zheng

## Abstract

Folding energy offers a useful metric for characterizing the stability and function of aptamers. However, experimentally measuring the folding energy is challenging and there is currently no general technique to measure this parameter directly. In this work, we present a simple approach for measuring aptamer folding energy. First, the aptamer is stretched under equilibrium conditions with a double-stranded DNA “molecular clamp” that is coupled to the aptamer ends. We then measure the total internal energy of stressed DNA molecules using time-lapse gel electrophoresis and compare the folding and unfolding behavior of molecular clamp-stressed molecules that incorporate either the aptamer or unstructured random single-stranded DNA in order to derive the aptamer folding energy. Using this approach, we measured a folding energy of 10.40 kJ/mol for the HD22 thrombin aptamer, which is consistent with other predictions and estimates. We also analyzed a simple hairpin structure, generating a folding energy result of 9.05 kJ/mol, consistent with the value predicted by computational models (9.24 kJ/mol). We believe our strategy offers an accessible and generalizable approach for obtaining such measurements with virtually any aptamer.

## Introduction

Aptamers offer the powerful capability to recognize and specifically bind to a wide range of biomolecules, including small molecules ^1^, metal ions ^2^, proteins ^3^, and even whole cells ^4^. An aptamer’s function is dependent upon its capacity to fold into and maintain a binding-competent conformation ^5^. As such, the folding energy for a given DNA or RNA sequence is an important determinant of aptamer stability, which in turn affects that aptamer’s target affinity and specificity, and thus its broader utility in molecular detection and other assays.

Unfortunately, it currently remains difficult to evaluate this parameter because of the difficulties associated with direct measurement of aptamer folding energy at equilibrium. One approach is based on time-dependent single-molecule pulling experiments, in which oligonucleotides are stretched by atomic force microscopy (AFM) or optical tweezers ^6–9^. These experiments drive the system away from equilibrium, where the magnitude of the rupture force is related to the pulling rate. As a consequence, one must use the relationship between free energies and irreversible work ^10–12^ described by Jarzynski’s equality ^13^ to calculate free energy from repeated measurements. However, some have challenged the validity of this approach, arguing that its underlying assumption—*i.e.*, the connection between the microscopic work performed by a time-dependent force with the corresponding Hamiltonian—is theoretically inconsistent ^14–16^. Other groups have used ultraviolet spectroscopy or circular dichroism (CD) to generate indirect measurements of melting curves based on distinct differences in the spectroscopic properties of folded and unfolded aptamers ^17,18^. However, the arbitrary determination of melting curve baselines can introduce considerable uncertainty, and the assumption that denaturation is only a two-state process with zero heat-capacity change greatly limits the universal application of this approach ^19,20^. Others have also performed melting experiments using differential scanning calorimetry (DSC),^18,20^ which measures differential heat capacity as a function of temperature ^19,20^. However, the accuracy of this approach is not guaranteed, because the ability to correctly determine the values of many thermodynamic parameters is heavily dependent on the choice of appropriate melting models ^20,21^.

We describe here a general strategy for directly measuring aptamer folding energy, which eliminates these confounders and sources of bias that can lead to inaccurate measurement. Our method employs a short piece of double-stranded (ds) DNA, which is shorter than the dsDNA persistence length of ~50 nm, as a “molecular clamp” to pull an aptamer into an unfolded state under equilibrium conditions. Once the appropriate length of the molecular clamp required for successful extension of the aptamer has been determined, we measure the total energy of the constrained molecule – including the bending energy of the molecular clamp and stretching energy of the aptamer – through a simple time-lapse gel electrophoresis method. The folding energy is calculated by comparing against the gel profile of a randomized, unstructured single-stranded (ss) DNA molecule that has been incubated with the same molecular clamp. Here, we demonstrate this method to determine the folding energy of the thrombin aptamer HD22 ^22^, and derive a measurement that is consistent with prior estimates. Our ‘folding energy measurement by molecular clamping’ (FE-MC) method only requires standard gel electrophoresis equipment and reagents, and therefore offers a simple and broadly accessible solution for characterizing aptamer folding and stability.

## Results and Discussion

### Overview of the FE-MC method

The central idea behind the FE-MC method is to exploit short stretches of dsDNA that act as “molecular clamps” to unfold aptamers. This requires the design of two DNA oligonucleotides: Strand A contains the aptamer sequence at the center, and is flanked by sequences that are complementary to Strand B (**Fig. 1A**, left). Therefore, hybridization between Strand A and Strand B produces a circularized, partially double-stranded molecule (**Fig. 1A**, right). This structure can be viewed as two coupled (nonlinear) springs – representing the dsDNA and ssDNA component – which are constrained at the same end-to-end distance (EED) *x*. This represents the distance between the two ends of the molecule after either bending for the dsDNA or stretching for the ssDNA, and describes the degree of deformation for both components. We have termed this circularized molecule a ‘stressed aptamer molecule’ (SAM), with a high internal energy that is stored in the bending energy of the dsDNA component (*E_d_*) and the stretching energy of the ssDNA component (*E_s_*). The dsDNA component applies stretching force at pico-Newton (pN) scales on the ssDNA component, *i.e.* the aptamer sequence, at equilibrium ^23^, the stretching behavior of which is modeled as shown in **Fig. S1A**. Since shorter molecular clamps would be expected to exert a stronger extending force (as explained in the Supplementary Information, SI), the aptamer may be successfully stretched into an unfolded state (**Fig. 1A**, lower right) or remain folded (**Fig. 1A**, upper right) depending on which state has lower internal energy for varying length of the dsDNA component (*N_d_*), as illustrated in **Fig. S1B**.

**Figure 1.**
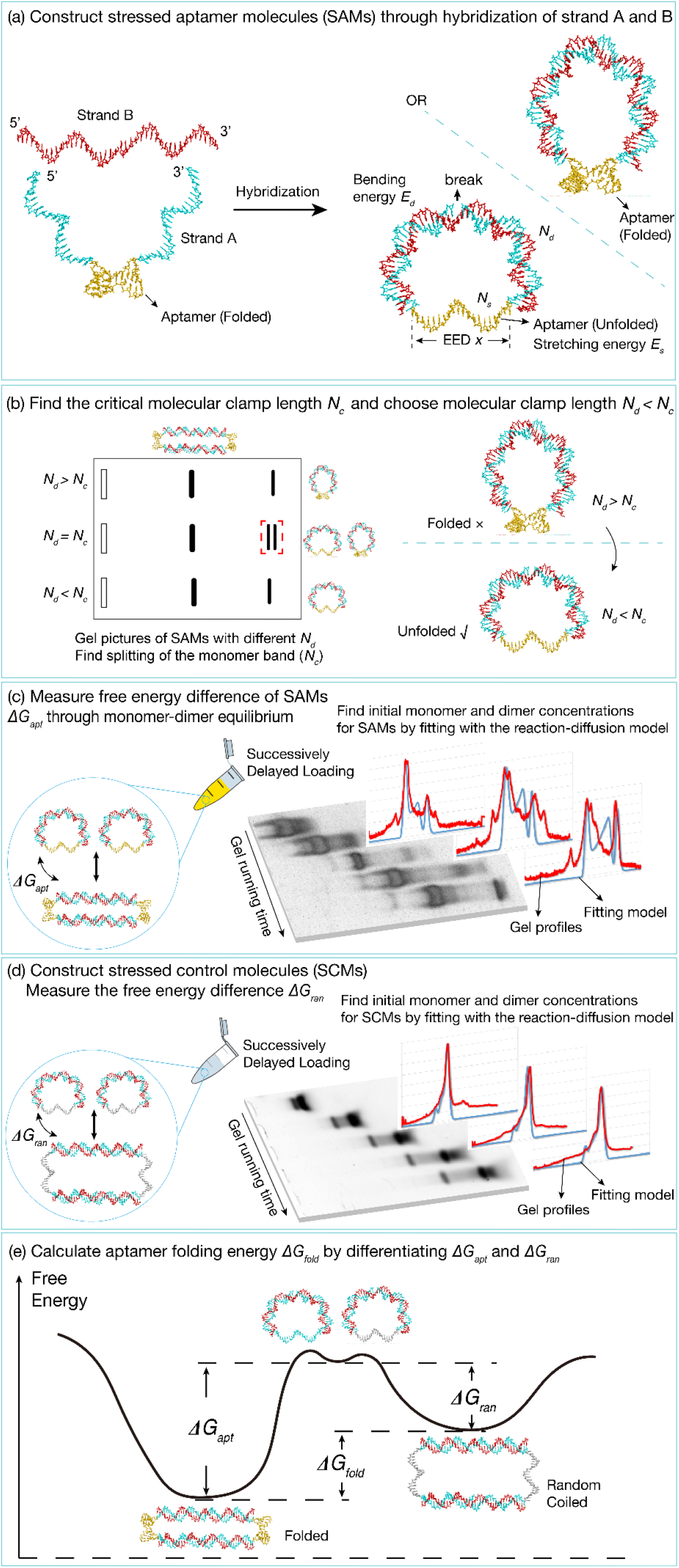
Determination of aptamer folding energy using the molecular clamp (FE-MC) method. (A) This procedure entails constructing a series of stressed aptamer molecules (SAMs). These comprise two strands: Strand A contains an aptamer sequence at its center (yellow), flanked by sequences (cyan) complementary to Strand B (red). Hybridization produces a circularized structure with a ‘break’ in one strand in the middle of the dsDNA component. The dsDNA component (with length *N*_*d*_) serves as an ‘molecular clamp’ that applies pN forces on the ssDNA aptamer component (with length *N*_*s*_). There is high internal energy stored in the dsDNA bending energy and ssDNA stretching energy, and depending on the length of *N*_*d*_, the aptamer may be stretched into an unfolded state (upper right) or remain folded (lower right). (B) The next step of FE-MC is to find the critical molecular clamp length (*N*_*c*_) at which the aptamer transitions from folded to unfolded. This entails gel electrophoresis of SAMs with different *N*_*d*_ values, yielding bands corresponding to monomers, dimers and other multimers arising from polymerization of multiple copies of Strand A and Strand B. *N*_*c*_ can be identified as the sample in which splitting of the monomer band is observed (red dashed box), indicating the coexistence of folded and unfolded aptamer states. A SAM in which *N*_*d*_ < *N*_*c*_ is chosen for subsequent steps. (C) We next measure the internal energy (*ΔG*_*apt*_) of the chosen SAM. Because of the break in the molecular clamp, this internal energy may be relaxed by forming dimers, in which the dsDNA component does not bend and the aptamer is not stretched. We can then quantify the SAM internal energy using Eq. (1) by measuring the equilibrium concentrations of monomers and dimers. Due to the concentration bias induced by ongoing monomer-dimer interconversion even during the gel electrophoresis, we obtain initial equilibrium concentrations by running time-lapse gel electrophoresis with successively delayed loading of multiple lanes with aliquots from the same SAM sample, and fitting ‘snapshots’ of gel profiles (red curves) with the reaction-diffusion model (blue curves). (D) In parallel, we construct stressed control molecules (SCMs) in which the aptamer from the SAM is substituted with an unstructured randomized sequence, and again measure the internal energy (*ΔG*_*ran*_). (E) This plot depicts the energy levels occupied by the monomeric and dimeric states of the SAM and SCM. Note that the energy level is the same for the unfolded SAM and SCM monomers, but the dimer energy levels are different because the aptamer is folded while the randomized sequence is unstructured. The aptamer folding energy *ΔG*_*fold*_ is calculated as *ΔG*_*apt*_ − *ΔG*_*ran*_.

The next step entails identification of the appropriate *N_d_* value to ensure that the aptamer is fully extended upon hybridization. To achieve this, we construct a series of SAMs with molecular clamps of varying *N_d_* and subject these constructs to gel electrophoresis. The resulting gel will show multiple bands corresponding to monomers, dimers, and other multimers arising from polymerization of multiple copies of Strand A and Strand B. In this step, we need to identify the value of *N_d_* at which splitting of the monomer band is observed (**Fig. 1B**), indicating the coexistence of two possible configurations with similar internal energy (as depicted in **Fig. S2**). This represents the transition length (*N_c_*) of the molecular clamp: when *N_d_* < *N_c_*, the aptamer is stretched and unfolded, whereas when *N_d_* > *N_c_*, the aptamer remains in a folded state.

We then select a SAM for which the molecular clamp length *N_d_* is smaller than *N_c_* for further analysis. It is important to note that there is a break in the middle of the dsDNA component of the SAM, which arises from the two ends of the aptamer-flanking ‘arms’ of Strand A after hybridization with Strand B. This break means that the SAM may relax its internal elastic energy through the formation of multimers, in which the dsDNA component is not forced to bend and the ssDNA component is not subject to stretching ^24^. Because polymerization lowers the entropy of molecules, there exists a chemical equilibrium of monomer-dimer interconversion, which can be observed through gel electrophoresis, with multiple bands corresponding to dimers and other multimers (**Fig. 1C**).

Quantitatively, this monomer-dimer interconversion can be described in terms of the equality of monomer and dimer chemical potentials, namely 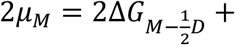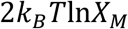 and *μ*_*D*_ = *k*_*B*_*T*ln*X*_*D*_. The internal energy of the SAM is hence described by the following equation:

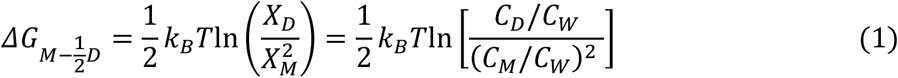

where *X_D_*, and *X_M_* are the molar fractions and *C_D_* and *C_M_* are the concentrations of the monomer and dimer, respectively. *C_W_* = 55 M is the concentration of water (see Methods for deriving details). Eq. (1) suggests that the internal energy of the SAM can simply be computed by quantification of monomer and dimer populations at equilibrium (*e.g.*, through gel electrophoresis). It is worth noting that this monomer-dimer interconversion continues even during gel electrophoresis, leading to bias in monomer and dimer concentrations relative to initial equilibrium values. To address this problem, we have adopted a ‘time-lapse’ gel electrophoresis method (**Fig. 1C**, right). Specifically, multiple aliquots of the same chosen SAM sample (in which *N_d_* < *N_c_*) are loaded into different lanes at successive times. The gel image for this SAM is then collected to show the evolution of monomer and dimer bands with time, enabling extrapolation back to initial equilibrium concentrations. A reaction-diffusion model is applied with the initial monomer and dimer concentrations as fitting parameters, and their values are determined so that the gel profiles at each running time are fitted to model curves with the same set of parameters (**Fig. 1C**, right; see Methods for details). The initial concentrations are then plugged into Eq. (1) to calculate the internal energy *ΔG*_*apt*_ within the SAM.

Next, we construct stressed control molecules (SCMs), in which the aptamer sequence in Strand A has been replaced with a randomized sequence (**Fig. 1D**). We extract the internal energy within the SCM (*ΔG*_*ran*_) in the same way as for the SAM. Finally, we compute the aptamer folding energy as follows: *ΔG*_*fold*_ = *ΔG*_*apt*_ − *ΔG*_*ran*_ (**Fig. 1E**). Since the aptamers in the SAM dimers are in a folded state while the randomized sequences in the SCM dimers are in a random-coiled state, *ΔG*_*apt*_ − *ΔG*_*ran*_ cancels the contribution of base-pairing in half of the SAM and SCM dimers, and yields the energy difference between the folded aptamer and the random DNA coil (see SI).

### Folding energy measurement of the HD22 thrombin aptamer

To demonstrate the FE-MC method, we chose an aptamer that recognizes human α-thrombin (HD22) ^22^. This 29-nt aptamer (5’-AGTCCGTGGTAGGGCAGGTTGGGGTGACT-3’) contains a duplex/G-quadruplex mixed structure ^25^, and exhibits strong target affinity (*K_D_* ~ 0.5 nM). We constructed a series of HD22 SAMs with *N_d_* ranging from 18–24 bp. Time-lapse gel images of these SAMs are shown in **Fig. 2**. When *N_d_* = 20 bp, splitting of the monomer bands was observed (**Fig. 2B**), indicating that this was the *N_c_* of the molecular clamp; when *N_d_* < 20, the HD22 aptamer was unfolded, whereas when *N_d_* > 20, the HD22 aptamer remained in the folded state.

**Figure 2.**
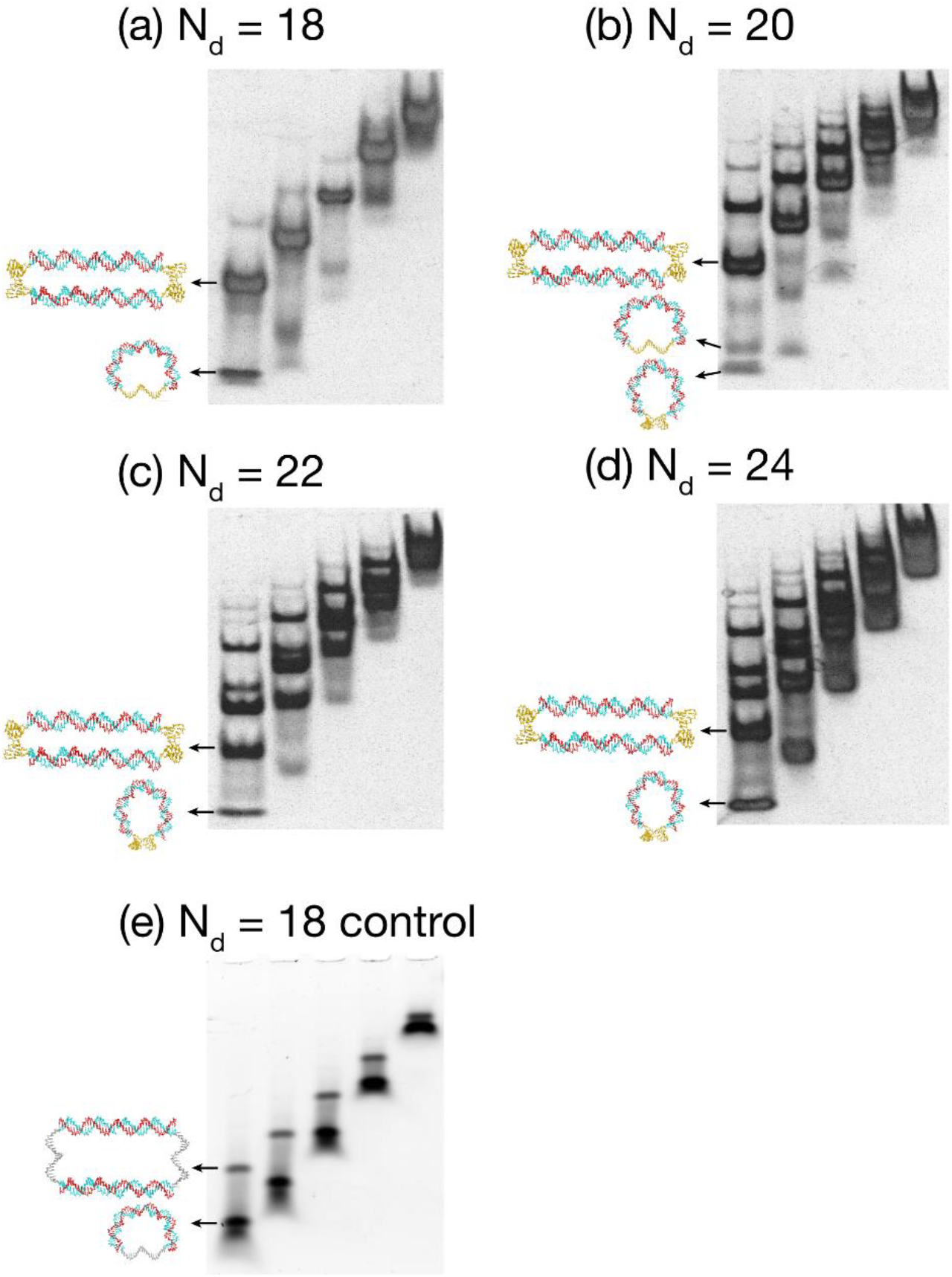
Time-lapse gel images for SAMs with the HD22 aptamer and *N*_*d*_ = A) 18, B) 20, C) 22, and D) 24 bp. (E) Time-lapse gel pictures for an SCM with a randomized control sequence and *N*_*d*_ = 18.

We selected an unfolded SAM (*N_d_* = 18 bp; SAM18) for subsequent detailed analysis. A time-lapse gel image of SAM18 was collected as shown in **Fig. 2A** and the gel profiles in different lanes (*i.e.*, at different running times) were successfully fitted with the reaction-diffusion model (**Fig. S3A**, see SI for fitting details). We found that the initial equilibrium concentrations of monomer and dimer were 0.42 μM and 0.79 μM, respectively. We hence calculated the internal energy of the SAM18 monomer 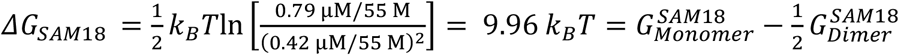 using Eq. (1). This describes the free energy difference between the SAM18 monomer 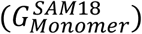 and half of the SAM18 dimer 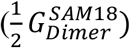.

We next constructed SCM18, with the same molecular clamp, but where Strand A incorporated a control 29-mer randomized sequence (5’-CAGCAGGCAATCGATACACACACAGTAGA-3’) with no secondary structure (checked by the DINAMelt web server ^26^). The time-lapse gel image was collected as shown in **Fig. 2E**. The internal molecular energy was calculated using Eq. (1) by fitting the SCM18 gel profiles with the reaction-diffusion model (**Fig. S3B**). We found that the initial monomer and dimer concentrations for the SCM were 1.78 μM and 0.11 μM, respectively (see SI for fitting details). Thus, 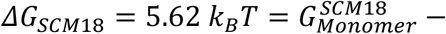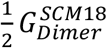.

SAM18 and SCM18 shared the same free energy in their stretched state, but the aptamers in the SAM18 dimers were folded while the randomized sequences in the SCM18 dimers were in a random-coiled state, leading to a difference in energy levels. Accordingly, the differential between *ΔG*_*SAM18*_ and *ΔG*_*SCM18*_ canceled the base-pairing energy in half of the SAM and SCM dimers, yielding the energy difference between the folded aptamer and the random DNA coil. Specifically, this can be calculated as follows:

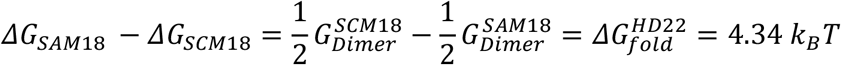

or 10.40 kJ/mol for the HD22 folding energy.

### Validation of the experimental results

In order to validate this measurement, we calculated the critical molecular clamp length (*N_c_*) to check whether it was consistent with experimental observations. The theoretical value of *N_c_* was determined for a scenario in which the folded and extended states of the SAM have similar energy values. Our determination of the folding energy for HD22 (4.34 *k*_*B*_*T*) was plugged into the model of the aptamer folding energy profile (see SI) as the ssDNA stretching energy of the SAM. The EED *x* between the two ends of the SAM was used to characterize the degree of deformation for the two SAM components. The total energy (*E_tot_*) of the SAM as a function of the EED *x* was calculated using the sum of the ssDNA stretching energy and dsDNA bending energy (see SI). We found that when *N_d_* = 20, the local energy minimum of the folded state was 4.95 k_B_T, while that of the stretched state was 5.20 k_B_T, as shown in **Fig. S2**. The similarity of the two energy values indicates that *N_d_* = 20 is the theoretical critical length, consistent with our experimental results.

We also compared the final HD22 folding energy value 4.34 k_B_T (or 10.40 kJ/mol) to other estimates. Prior studies of aptamer folding based on pulling experiments using AFM or optical tweezers have yielded free energy estimates for the complete unfolding of the *pbuE* riboswitch aptamer in the range of 2–3 kcal/mol (8.4–12.6 kJ/mol) ^9^, the same order of magnitude for energy scale as we obtained with HD22 using our orthogonal method. For this particular aptamer, the DINAMelt web server ^26^ predicts a folding energy of just 1.11 kcal/mol (4.67 kJ/mol), which is much smaller than the value we obtained. This is because the DINAMelt algorithm does not account for the energy of the 3D structure of the G-quadruplex. According to previous studies using UV-spec or DSC, the ultimate free energy *ΔG*(310 K) for a low-order G-quadruplex is approximately 1.50 kcal/mol (6.30 kJ/mol) ^17,19^. Since HD22 features a combination of duplex and G-quadruplex structure ^25^, a rough estimate from adding the base-pairing energy of the duplex (4.67 kJ/mol) and the G-quadruplex folding energy (6.30 kJ/mol) yields a total folding energy of 10.97 kJ/mol for HD22, which is very similar to the value we calculated (10.40 kJ/mol).

Such an estimate is necessary given the lack of directly comparable, precise folding energy predictions by other metrics, but we were also able to confirm the accuracy of our FE-MC approach with a simple hairpin structure (5’-TTGTCAT TTTTTTTTTTTTTTTATGACTT-3’). The folding energy of this simple structure is easily predicted by DINAMelt ^26^ to be 9.24 kJ/mol under ambient conditions, and FE-MC yielded a highly consistent measurement of 9.05 kJ/mol (see SI for details), further supporting the validity and accuracy of our approach.

## 4. Conclusion

In this work, we describe a simple approach for probing the folding energy of aptamers. In our ME-FC method, the aptamer is unfolded by a dsDNA component that acts as a molecular clamp under equilibrium conditions. The internal energy of this stressed aptamer molecule is determined using time-lapse gel electrophoresis, and then compared against a similar stressed DNA molecule in which the aptamer has been substituted with a randomized ssDNA with no secondary structure. The internal energy difference between the two stressed molecules yields the aptamer folding energy.

Using this approach with the HD22 thrombin aptamer as a model, we computed a folding energy of 4.34 k_B_T (~10.4 kJ/mol), which is in keeping with aptamer folding energy scales from other studies as well as rough predictions based on HD22’s structure. We also validated the accuracy of FE-MC by correctly measuring the folding energy of a hairpin structure, obtaining results that mirror the value predicted by the DINAMelt web server. Although we have calculated the folding energy measurement of a relatively short aptamer (29 nt) here, FE-MC method should also be applicable to aptamers with longer lengths of 60–80 nt. This is valuable in that many newly-selected aptamers include primer-biding sites and have not been minimized. Since FE-MC only requires time-lapse gel electrophoresis experiments, we believe it offers a highly accessible and generalizable method for evaluating aptamers in a simple and direct manner.

## Methods and Materials

### Sample preparation

All oligonucleotide sequences (listed in **Table 1**) were synthesized by Sangon Biotech (Shanghai) with HPLC purification. The red sequences of Strand A hybridize with the corresponding Strand B. We confirmed hybridization of these dsDNA regions by checking sequence thermodynamic properties with the DINAMelt web server ^26,27^.

**Table 1.**
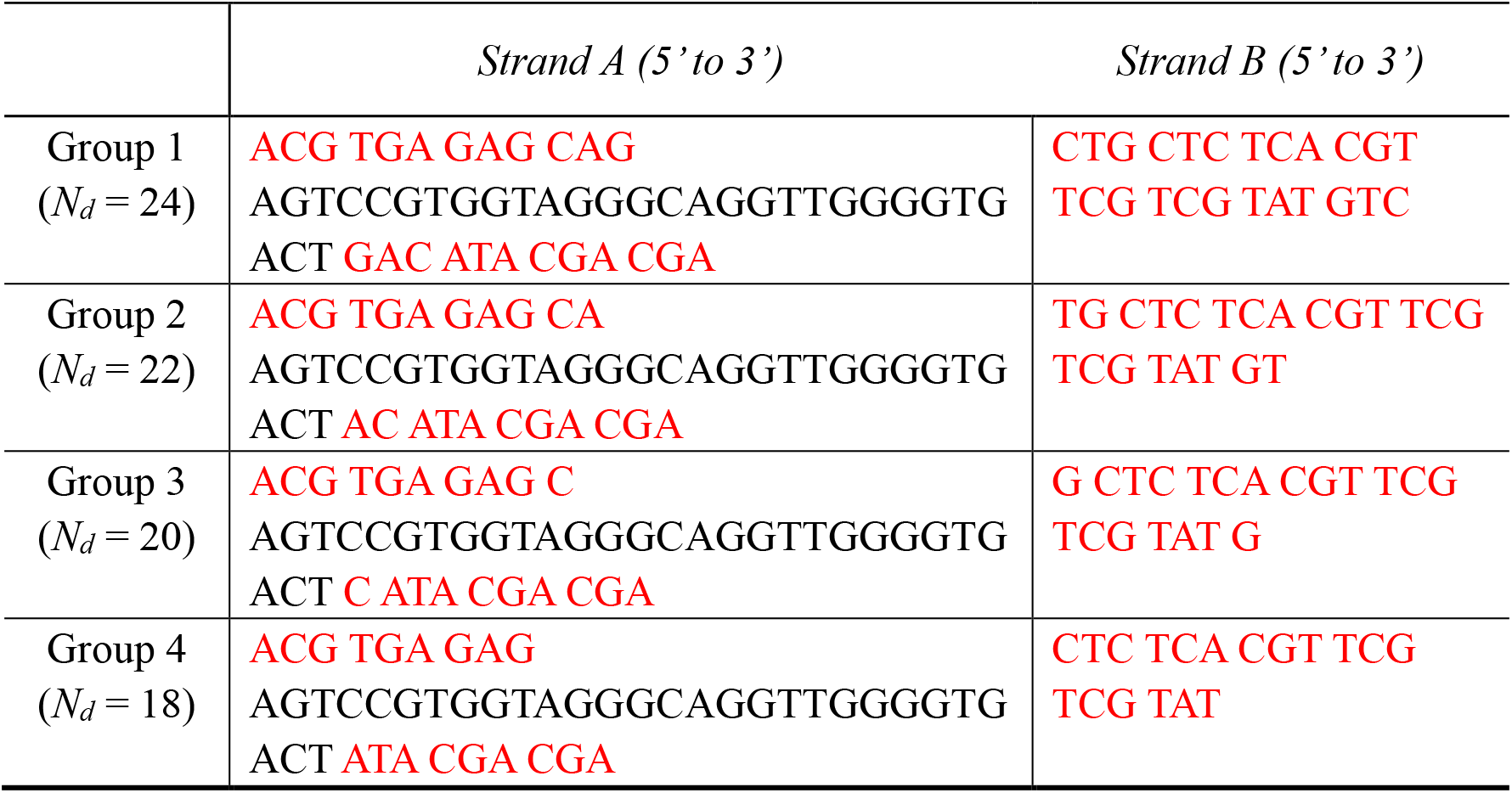
DNA sequences used to form SAMs with different *N*_*d*_.

Each Strand A was mixed with equimolar amounts of Strand B and diluted to a final concentration of 2 μM and a final volume of 200 μL in the hybridizing buffer (10 mM Tris, 5 mM MgCl_2_, 100 mM NaCl, pH = 7.9). The mixture was then heated at 95 ℃ for 10 min and annealed at room temperature overnight (~0.065 ℃/min) to ensure the proper conformational structures in stressed DNA molecules.

### Molecular energy extraction based on monomer-dimer equilibrium

At equilibrium of monomer-dimer interconversion, the chemical potentials for two monomers 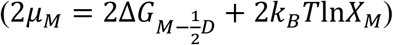 and one dimer (*μ*_*D*_ = *k*_*B*_*T*ln*X*_*D*_) are equal: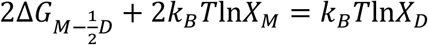. Since the base-pairing is identical for one dimer and two monomers, the energetic contribution of base-pairing cancels out and the internal elastic energy stored in the monomer can be characterized by the following equation: ^24^

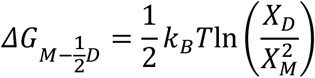

where *X*_*D*_ = *C*_*D*_/*C*_*W*_ and *X*_*M*_ = *C*_*M*_/*C*_*W*_, describing the molar fraction of the monomer and dimer respectively, and *C*_*M*_ and *C*_*D*_ are the respective concentration of the two species. *C*_*W*_ = 55 M is the concentration of water. This equation essentially gives the energy difference between the monomer and half of the dimer (as the ground state), and suggests that this energy difference can be simply extracted by measuring the equilibrium monomer and dimer concentrations.

### Time-lapse gel electrophoresis

Concentrations of monomers and dimers were experimentally determined by time-lapse gel electrophoresis with a 5% polyacrylamide gel, based on the intensities of the corresponding gel bands (using necessary calibrations as described in ^24^). The monomer-dimer interconversion of the stressed DNA molecules also occurs while the sample is moving through the gel and has to be taken into account. Therefore, in order to extrapolate back the initial equilibrium concentrations of the population of monomers and dimers at *t_0_*, we loaded different lanes of the gel with aliquots from the same sample at intervals of 10 min to capture “snapshot” gel profiles at different times. These various aliquots were run under 100 V for 55 min, 45 min, 35 min, 25 min, and 15 min, respectively. Images from time-lapse gel electrophoresis were converted to gel profiles using ImageJ ver1.51.

### Reaction-diffusion model

In order to extract the initial population of monomers and dimers, we fitted these gel profiles with a reaction-diffusion model ^28^ that has been successfully applied to the analysis of electrophoresis with protein-DNA ^29^, hairpin-duplex ^30^, and monomer-dimer ^23,24^ interconversions. The model formulas are:

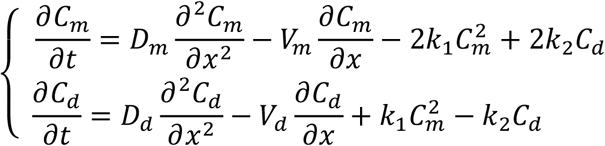

where *C*_*m*_ and *C*_*d*_ are the concentration of monomers and dimers respectively, as a function of gel running time *t* and gel vertical position *x* from the bottom of the loading well; *D*_*m*_ and *D*_*d*_ are the diffusion constant of the two species, and *V*_*m*_ and *V*_*d*_ are their mobilities; *k*_*1*_ is the conversion rate from monomers to dimers, while *k*_*2*_ is the rate for the reverse reaction. During fitting with the reaction-diffusion model, *V*_*m*_ and *V*_*d*_ were first determined by measuring the distance that the population had shifted at various running times. *D*_*m*_ and *D*_*d*_ were determined from the width of the corresponding peaks of monomers and dimers. The initial monomer and dimer concentrations (*C*_*m0*_ and *C*_*d0*_) and interconversion rates (*k*_*1*_ and *k*_*2*_) were adjusted so that the model could fit the gel profiles at different times (*i.e.*, in different lanes) with the same set of parameters. Finally, *C*_*m0*_ and *C*_*d0*_ were used to calculate the internal energy of the stressed DNA molecules.

Note that the computed elastic energy was rather insensitive to the parameter values in the model. Even a barely fair fitting to the gel profiles, in which the interconversion rates were set to zero, leads to a difference of ~15% in the initial concentration values, which in turn resulted in only a ~0.2 k_B_T change in the final elastic energy result due to the logarithmic function in Eq. (1).

## Acknowledgements

This study was supported by the National Key R&D Program of China (2017YFC1600603), the National Natural Science Foundation of China (21705031), the Natural Science Foundation of Anhui Province (1808085QB39), the Fundamental Research Funds for the Central Universities (PA2019GDQT0018), and Engineering Research Center of Bio-process by Ministry of Education in Hefei University of Technology. This work was also supported by the Chan-Zuckerberg Biohub (HTS).

## Supplementary Information

### Use of dsDNA as molecular clamp

Long chains of dsDNA normally exhibit bending elasticity resembling an entropic spring, in accordance with the ‘wormlike chain’ model (WLC) with energy of:

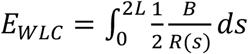

where *R* is the radius of curvature, *s* the arc length along the rod, *B* the bending modulus (*B* ≈ 200 pN × nm^2^), and *2L* the contour length (*N*_*d*_ × 0.33 nm) ^31,32^. However, this model breaks down when the contour length is less than the persistence length of dsDNA (*i.e.*, 150 base pairs, or 50 nm). We have previously determined that this breakdown involves a local “kink” forming in the middle of the dsDNA, which may greatly lower the bending energy of the molecule after a critical deformation. The following is an analytical expression of the dsDNA bending energy (*E*_*d*_) at this length:

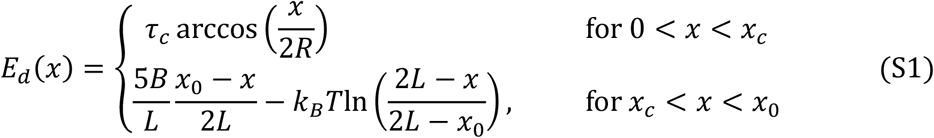

where τ_c_ is the critical torque at which the kink develops; *x* the end-to-end distance (EED); *R* = *L*(1−2*γ*^2^/45) is the arc length, where *γ* = *Lτ_c_*/(2*B*); *x*_0_ = 2*L*[1−*k*_*B*_*TL*/(5*B*)] is the EED at zero force; and *x_c_* is the critical EED ^23,24,33^. The lower form of the equation corresponds to the smoothly bent solution, while the upper describes the kinked state. Therefore, the bending behavior – including the applicable force for using short dsDNA as molecular clamp – is comprehensively known. For example, according to Eq. (S1), the bending elastic energy of dsDNA shows an increase with decreasing length (*N*_*d*_). Hence, the stretching state of the aptamer can be readily tuned by adjusting *N*_*d*_.

### Model of aptamer folding energy profile

The stretching force applied to the aptamer in the context of the clamp initially leads to disruption and unfolding of its structure, with a corresponding free energy *ΔG*_*fold*_. After the aptamer is completely unfolded at a critical deformation, it returns to a coiled state as simple ssDNA without any internal structures, and the free energy required for pulling is purely elastic, arising from the entropy change of the ssDNA as a random coil (*ΔG*_*coil*_). The random coil state follows the WLC model described by the polynomial expansion of the Marko-Siggia expression ^7^:

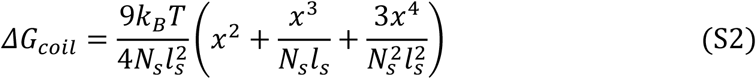

where *x* is the EED, *N_s_* the number of bases of ssDNA, and *l_s_* the persistence length of ssDNA (~0.7 nm). Therefore, the stretching energy for the aptamer can be modeled as the superposition of an energy well with a depth of *ΔG*_*fold*_ in the elastic energy of random coil *ΔG*_*coil*_, as described in Eq. (S2) ^31^ and shown in **Fig. S1A**. The detailed energy landscape of aptamer folding energy is not discussed in this paper.

### The effect of molecular clamps on aptamer folding states

For FE-MC to succeed, the dsDNA molecular clamp must be capable of pulling the aptamer apart from the folded state. To understand the effect of molecular clamps on aptamer folding states, we have calculated the total energy profile using the energy model for stretching energy of the ssDNA component (**Fig. S1A**), with the depth of the energy well of the experimentally measured HD22 aptamer folding energy (4.34 k_B_T), and Eq. (S1) for bending energy of the dsDNA component. We found that aptamers situated within stressed DNA molecules may shift from a folded to an extended state as the length of the molecular clamp decreases, producing “stiffer” clamps. As simulated in **Fig. S1B**, there are two local minimums for the total energy vs EED for stressed DNA molecules containing the aptamer, corresponding to the stretched and folded states respectively. When *N*_*d*_ = 22, the local minimum with a small separation (*i.e.*, EED = 1.02 nm) is lower, and the energy partition is *E*_*d*_ = 9.39 k_B_T in the molecular clamp and *E_s_* = −4.19 k_B_T in the aptamer, where the majority of the energy lies in the molecular clamp. But when *N*_*d*_ = 18, the local minimum with a large separation (*i.e.* EED = 5.92 nm) becomes lower, and the energy partition is *E*_*d*_ = 0.01 k_B_T in the molecular clamp and *E_s_* = 4.76 k_B_T in the aptamer. In this context, the majority of the energy is stored in the aptamer, leading to its full extension.

**Figure S1.**
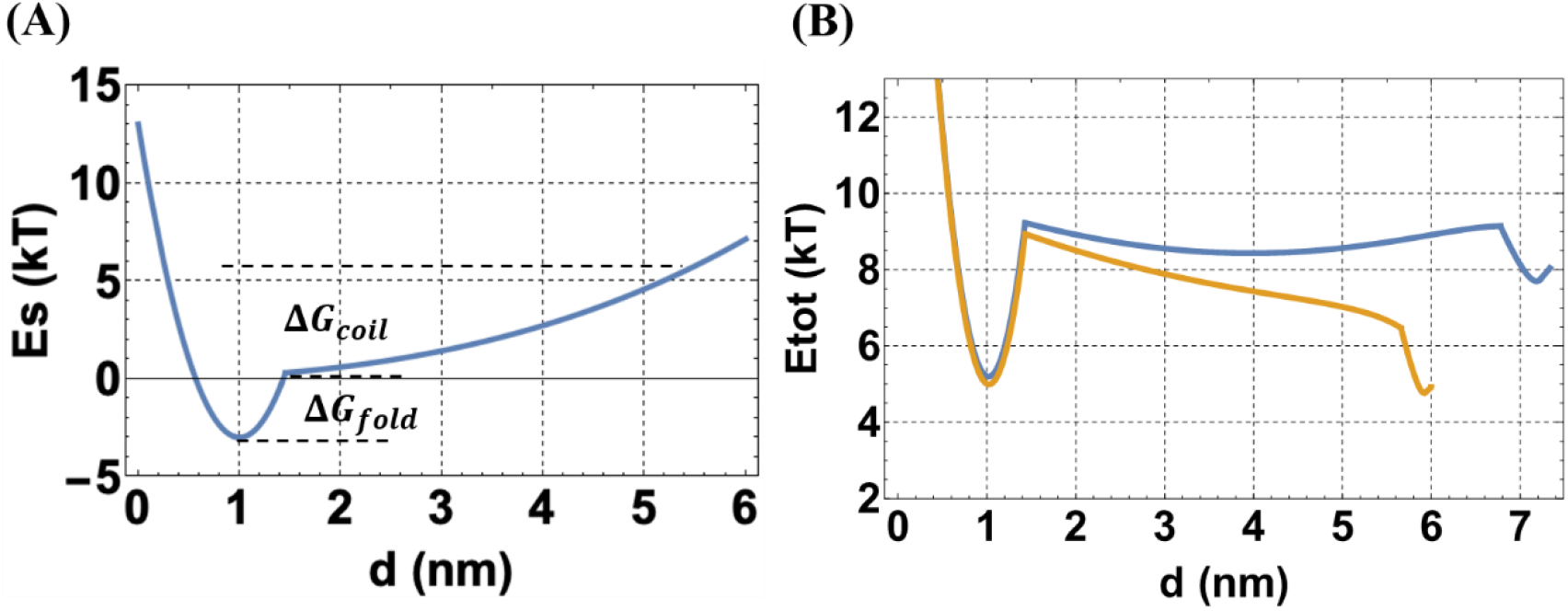
(A) The energy model for aptamer stretching: the superposition of an energy well with depth of *ΔG*_*fold*_ on the WLC energy *ΔG*_*coil*_ as expressed by Eq. (S2). (B) Calculated total energy profile (using Eq. (S1) and the aptamer energy model in A) *vs* EED for stressed DNA molecules with the HD22 aptamer and a molecular clamp of *N*_*d*_ = 22 (blue) or 18 (orange). The global energy minimum shifts from the local minimum on the left to the one on the right as *N*_*d*_ changes from 22 to 18, corresponding to the transition from the compressed state to the stretched state.

### Comparison of the experimental results with a theoretical model

For validation, the folding energy value obtained for the HD22 aptamer with FE-MC (4.34 k_B_T) was plugged into the theoretical model for energy profiles of SAMs to check whether it would produce the same critical molecular clamp length (*N_c_*) as we observed in the time-lapse gel. The total energy of the SAM with the HD22 aptamer as a function of the EED *x* (*E*_*tot*_ *vs x*) was calculated by taking the sum of the dsDNA bending energy (Eq. (S1)) and ssDNA stretching energy (see aptamer folding energy profile model above). The total energy profile for the SAM with *N*_*d*_ = 20 is shown in **Fig. S2**. The two local energy minimums correspond to the folded and stretched states, with energy values of 4.95 k_B_T and 5.20 k_B_T, respectively. This clearly indicates that the critical molecular clamp length *N*_*c*_ = 20, is consistent with our measurements after varying *N*_*d*_ as shown in **Fig. 2**, confirming the validity of the FE-SM method.

**Figure S2.**
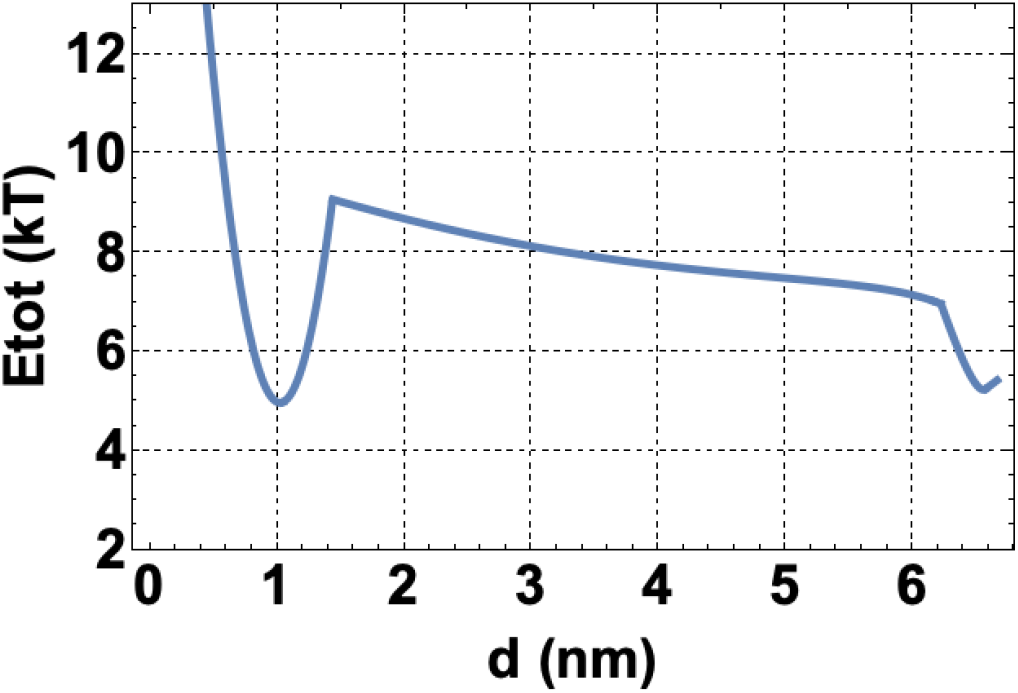
Calculated total energy profile (using Eq. (S1) and the aptamer energy model) *vs* EED for the SAM with the HD22 aptamer and a molecular clamp of *N*_*d*_ = 20. The measured HD22 folding energy value 4.34 k_B_T was plugged into the aptamer energy model for energy well depth. The two local energy minima corresponding to the folded and stretched states were calculated to be 4.95 k_B_T and 5.20 k_B_T, respectively. The similarity of these two values indicates that the critical length *N_c_* = 20, consistent with the experimental results shown in **Fig. 2**.

### Fitting details using the reaction-diffusion model

As described in Methods, the reaction-diffusion model was used for analysis of the time-lapse gel profiles for SAM18 and SCM18. For the SAM18 gel profiles, the vertical positions of the monomer and dimer bands (from the bottom of the loading well) at different running time were determined so that the mobilities for both species could be extracted using a linear fitting. We calculated *V*_*m*_ = 10.26 × 10^−2^ cm/min and *V*_*d*_ = 7.59 × 10^−2^ cm/min, respectively. The diffusion constants *D*_*m*_ and *D*_*d*_ were set according to the width of the corresponding peaks in the gel profiles and found to be *D*_*m*_ = 2.02 × 10^−5^cm^2^/min and *D*_*d*_ = 1.95 × 10^−5^cm^2^/min. The other parameters were adjusted so that the model might fit the experimental gel profiles at different times with identical values for these parameters. During the actual fitting, the heights of the monomer and dimer peaks were the key fitting factors. It should be noted that multiple sets of parameters may produce comparable fitting results for the gel profile at a given running time, but only a unique combination of parameters would work for all running times. For the gel profile of SAM18, *C*_*m0*_ = 0.42 μM, *C*_*d0*_ = 0.79 μM, *k*_*1*_ = 0.0001 μM^−1^ min^−1^ (monomer association rate), and *k*_*2*_ = 0.0249 min^−1^ (dimer dissociation rate) generated good fitting results for the gel profiles at 55, 45, and 35 min, as shown in **Fig. S3A**.

Gel profile analysis for SCM18 was slightly more complicated because there were two monomer bands in the gel patterns. Therefore, besides the interconversion between dimers and monomers, the interconversion between two monomer states had also to be considered. Following the same fitting procedures, the mobilities for the two monomer species and dimers were obtained depending on the vertical positions of the corresponding bands (*V*_*m*1_ = 6.87 × 10^−2^ cm/min, 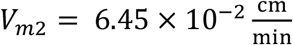, and *V*_*d*_ = 5.43 × 10^−2^ cm/min). The diffusion constants were identified to be *D*_*m*1_ = 1.95 × 10^−5^cm^2^/min, *D*_*m*2_ = 1.95 × 10^−5^cm^2^/min, and *D*_*d*_ = 1.37 × 10^−5^cm^2^/min. By adjusting the other fitting parameters, we found that the combination of *C*_*m10*_ = 1.63 μM, *C*_*m20*_ = 0.15 μM, *C*_*d0*_ = 0.11 μM, *k*_*1*_ = 0.0040 μM^−1^ min^−1^ (monomer association rate), *k*_*2*_ = 0.0027 min^−1^ (dimer dissociation rate), and *k_3_* = 0.0100 min^−1^ (monomer interconversion rate) would produce the best fitting results for the gel profiles at 55, 45, and 35 min, as shown in **Fig. S3B**.

**Figure S3.**
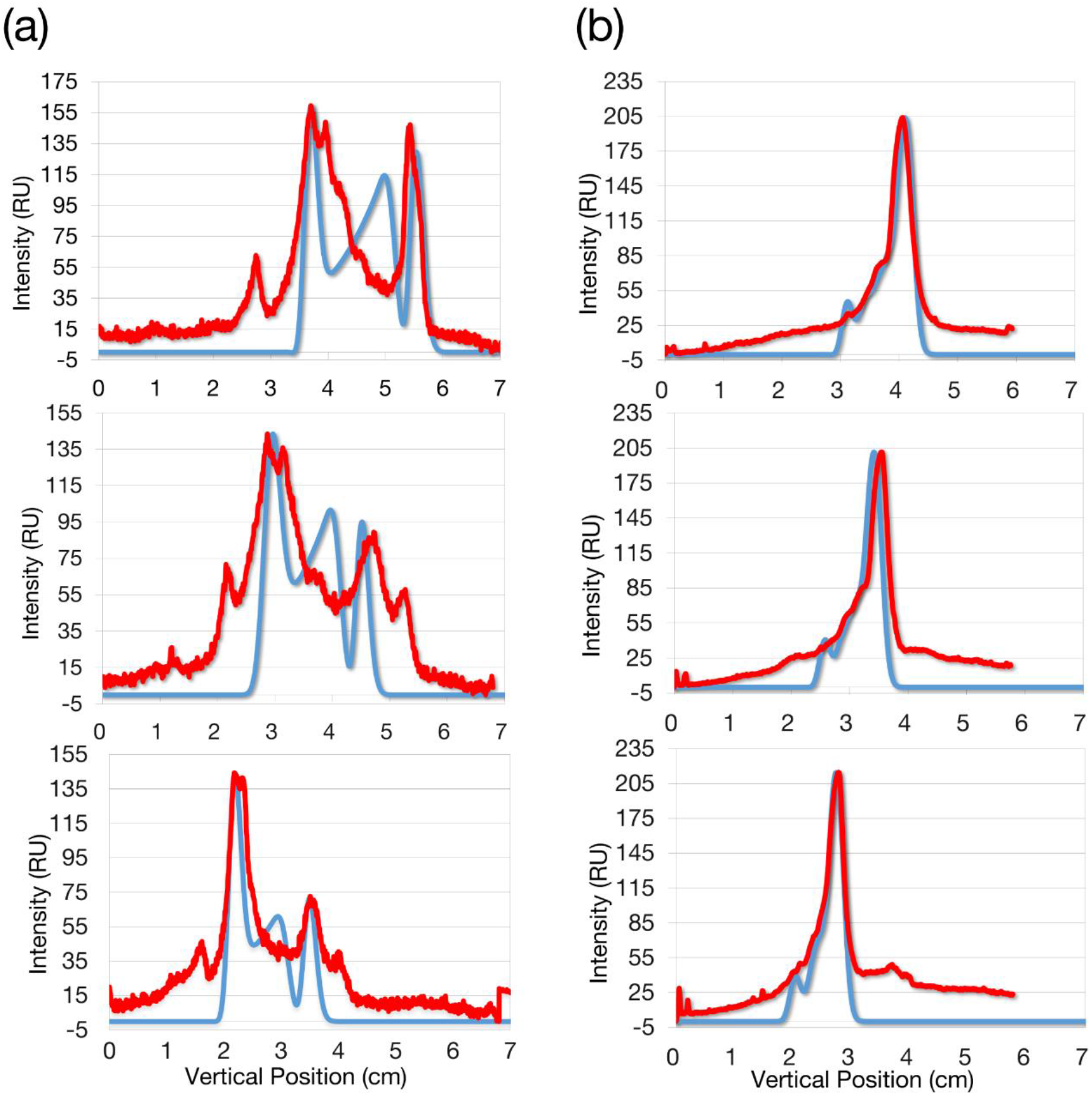
Intensity profiles of the 55, 45, and 35 min lanes from top to bottom of the time-lapse gel (red), and the fit with the reaction-diffusion model (blue) used to extract the equilibrium or initial values of the concentrations of the monomer and dimer for (A) SAM18 and (B) SCM18.

### Demonstration of the FE-MC method with hairpin structures

We also applied the FE-MC method to a hairpin structures (5’-TTGTCAT TTTTTTTTTTTTTTTATGACTT-3’)for which the folding energy has been well predicted by DNA folding algorithms ^26,27^. The hairpin had a total length of 29 nt, like HD22, and contained 5 bp in the stem and 15 nt in the loop. The folding energy predicted by the DINAMelt web server ^26^ was 2.16 kcal/mol (or 9.24 kJ/mol) under ambient conditions.

**Table S1.**
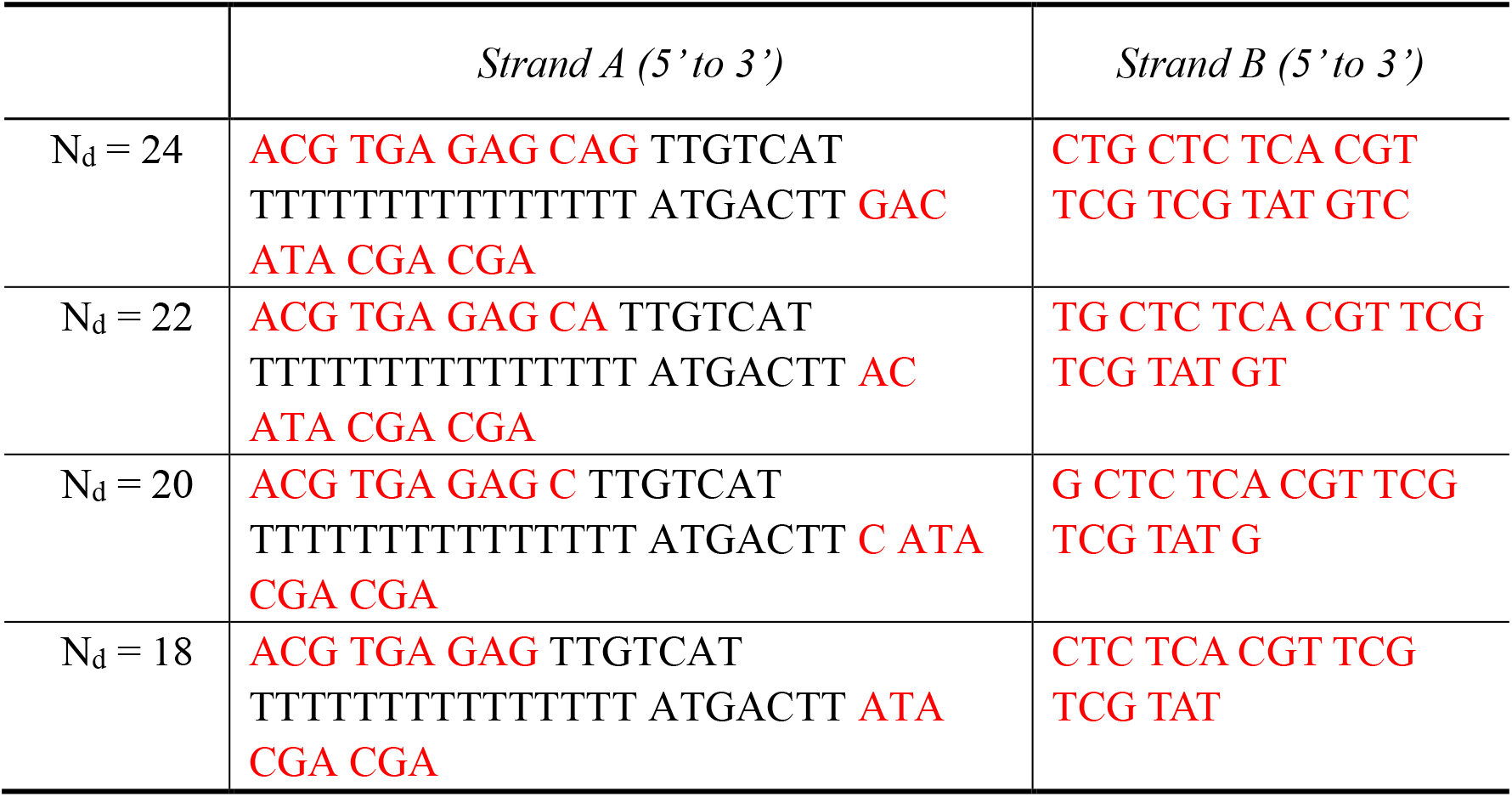
DNA sequences used to form hairpin SAMs with *N*_*d*_ ranging from 18–24 bp.

A series of SAMs containing this hairpin were prepared using the DNA sequences shown in **Table S1**, following the sample preparation procedures in Methods. Time-lapse gel images of these SAMs in each group were taken as shown in **Fig. S4A**. Splitting of the monomer bands was observed for the SAM with *N*_*d*_ = 20, indicating that this was the critical molecular clamp length. The SAM with *N*_*d*_ = 18 (denoted as ‘SHM18’) was thus chosen for subsequent detailed analysis. Again, we obtained the initial equilibrium monomer and dimer concentrations by fitting the time-lapse gel profiles of SHM18 (as shown in **Fig. S4**) using the reaction-diffusion model. The mobilities for both monomers and dimers were extracted using a linear fitting for vertical positions of the corresponding bands as a function of time and found to be *V*_*m*_ = 9.85 × 10^−2^ cm/min and *V*_*d*_ = 7.86 × 10^−2^ cm/min. *D*_*m*_ and *D*_*d*_ were found to be 2.0 × 10^−5^cm^2^/min and 1.9 × 10^−5^cm^2^/min, respectively. We found that *C*_*m0*_ = 0.53 μM, *C*_*d0*_ = 0.735 μM, *k*_*1*_ = 0.0005 μM^−1^ min^−1^, and *k*_*2*_ = 0.0190 min^−1^ generated the best fitting results for the gel profiles at 55, 45, and 35 min, as shown in **Fig. S4B**. The internal energy of the SHM18 was computed using Eq. (1) to be *ΔG*_*SHM18*_ = 9.39 *k*_*B*_*T*.

Since the total length of the chosen hairpin was 29 nt, we used the same SCM18 as in the main text as a control. We measured an internal energy for SCM18 of *ΔG*_*SHM18*_ = 5.62 *K*_*B*_*T*. Therefore, the folding energy of the hairpin could be computed by 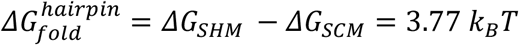, or 9.05 kJ/mol, which was highly consistent with the value predicted by the DINAMelt web server.

**Figure S4.**
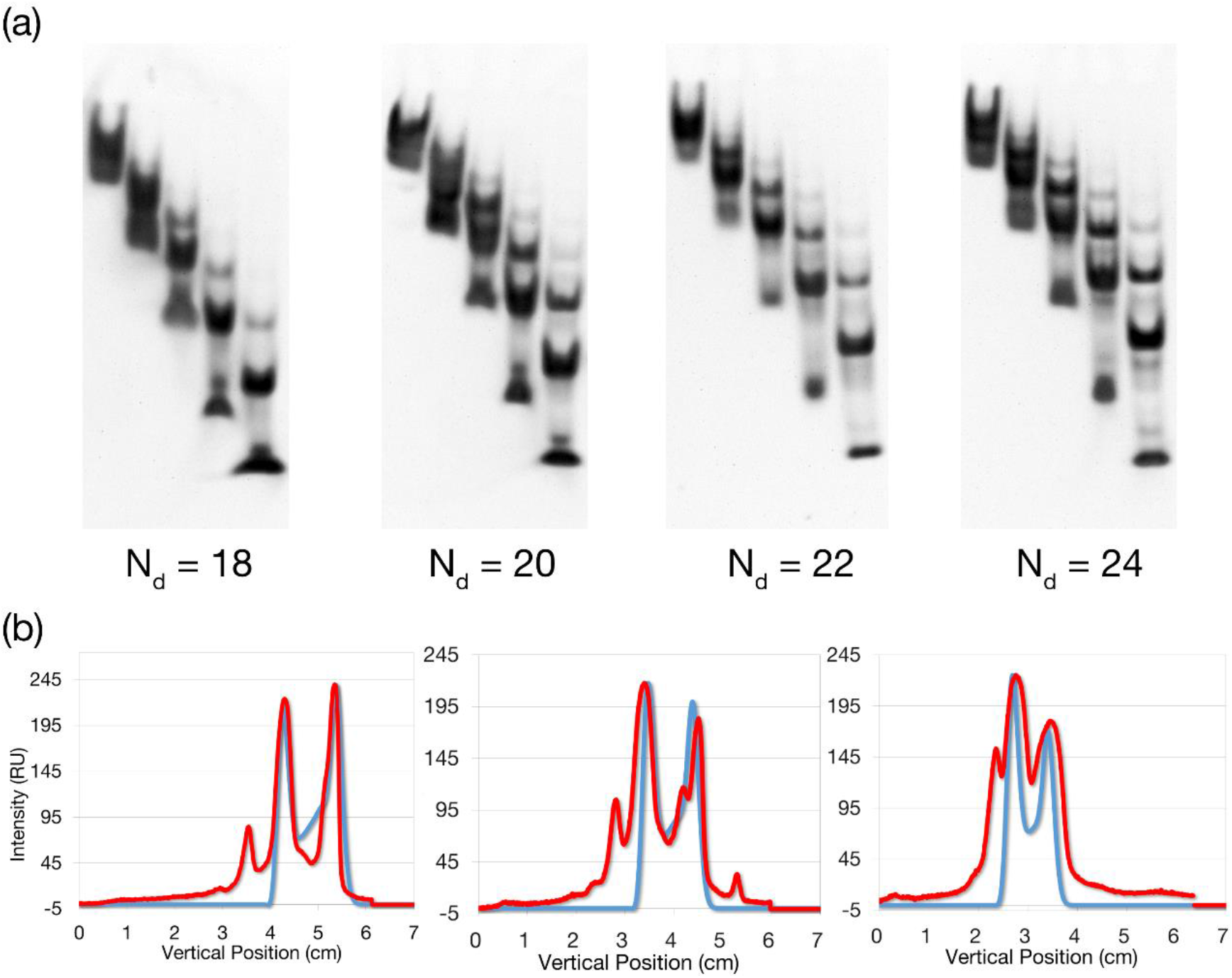
(A) Time-lapse gel pictures for SAMs with molecular clamps of *N*_*d*_ = 18, 20, 22, and 24. (B) Intensity profiles of the 55, 45, and 35 min lanes (from left to right) of the time-lapse gel (red), and the fit with the reaction-diffusion model (blue) used to extract the equilibrium or initial values of the concentrations of the monomer and dimer for the hairpin and the molecular clamp of *N*_*d*_ = 18.

